# Semi-quantitative detection for L-asparaginase producing fungi and the impact of carbon and nitrogen sources on enzyme activity

**DOI:** 10.1101/2021.02.02.429175

**Authors:** Mahdi Hamed, Ahmed A Osman, Mustafa Ateş

## Abstract

**Objective:** To semi-quantitively screen filamentous fungi isolated from different habitats for L-asparaginase production by three indicators; phenol red, cresol red and bromothymol blue and to examine the impact of different carbon and nitrogen sources on the enzyme production using different fungal isolates.

**Materials and methods:** Fifty-five fungal isolates were tested for L-asparaginase production by plate assay using Modified Czapek-Dox (MCD) medium. The enzyme activity was estimated using the Nessler method which measures the concentration of ammonia formed owing to the enzyme action on the substrate. The impact of nitrogen and carbon sources on the enzyme production was done by using the best three L-asparaginase producers from the semi-quantitative screening.

**Results and conclusions:** A total of 53/55 (96.36%) fungal isolates were L-asparaginase producing strains, of them, *Cladosporium tenuissimum, Penicillium camembertii* and *Aspergillus carneus* showed high enzyme production. Production of L-asparaginase was higher with the glucose and urea as carbon and nitrogen sources, respectively. The highest enzyme level (5,558 U/ml) was produced by *C. tenuissimum* in a glucose-containing medium. This study shows that *P. camemberti, A. carneus*, and *C. tenuissimum* are good L-asparaginase producers and thus could be used for L-asparaginase production

## Introduction

L-Asparagine amido hydrolase (L-asparaginase, E.C.3.5.1.1), is an enzyme catalyzes the conversion of L-asparagine to L-aspartic acid and ammonia [1]. L-asparaginase produced by plants, animals and microbes, and It has been used as effective treatment of children with acute lymphoblastic leukemia and in processing fried and baked food products which might contain a carcinogenic compounds[2].

Due to their ability to be cultured in an inexpensive substrate, easiness of optimization and purification techniques and to metabolically engineered, microbes are the preferred source of the enzyme [2, 3], however, pharmaceutically L-asparaginase is obtained primarily from *Escherichia coli* and *Erwinia carotova* is associated with undesired hypersensitivity and decreased enzyme activity [4, 5]. Thus, owing to the non-toxicity of L-asparaginase of an eukaryotic sources, filamentous fungi have been explored for the enzyme with least unfavorable effects [18].

## Material and methods

### Microorganisms

Fifty-five filamentous fungal isolates were obtained from the stocks of the Laboratories of three Turkish universities, Ege university, Adnan Mendres university and Celal Bayar university. Fungal isolates were maintained by cultivation in a slant of potato dextrose agar.

### Semi-quantitative production of L-asparaginase

Fungal strains screened for enzyme production on MCD medium, components g/litre; glucose: 2.0, L-asparagine: 10, KH_2_PO_4_: 3.0, Na_2_HPO_4_: 6.0, MgSO4.7 H_2_O: 0.50, NaCl : 0.50, FeSO_4_: 0.03, CuNO_3_. 3H_2_O 0.03, ZnSO_4_.7H_2_O 0.05, CaCl_2_: 0.05, agar: 20, indicator: 0.009) [6]. Inoculated plates were inoculated for 96 h at 30 °C. The strains that showed a zone of change in colour around the colonies indicated L-asparaginase production. Semi-quantitative screening was performed as described by Gonçalves et al. [7].

### L-asparaginase production via submerged fermentation

Fifty ml of sterile MCD production medium, g/litre (glucose: 2.0, L-asparagine: 10, KH_2_PO_4_: 1.52, MgSO4.7 H_2_O: 0.52, KCl : 0.52, FeSO_4_: 0.03, CuNO_3_. 3H_2_O 0.03, ZnSO_4_.7H_2_O 0.05) in a 250 ml Erlenmeyer flask was used for production. The bottles were incubated for 96 hours (30 °C and 150 rpm), then to separate the fungal cell mass, centrifuged at 5000 rpm for 15 minutes at 4 °C. The supernatant was used as a raw enzyme source to determine enzyme activity [8, 9, 10].

### Effect of carbon source

Using three isolates, *A. carneus, P. camemmberti and C. tenuissimum* the effect of glucose, starch, lactose, glycerol and sucrose on enzyme production was investigated. The MCD production medium was prepared with a concentration of 0.2% of the carbon source mentioned above. The flaks were incubated as mentioned above (section 2.3). Enzyme activity test was done after 96 hours.

### Effect of nitrogen source

Using three isolates (*A. carneus, P. camemmberti and C. tenuissimum*), the impact of proline, urea, asparagine, ammonium chloride and sodium nitrate on enzyme production was investigated. The production medium was prepared with a concentration of 0.5% of the nitrogen source mentioned above. 0.5% asparagine was added as an inducer for _L_-asparaginase production. The flasks were placed as in section 2.3. Enzyme activity test was done after 96 hours.

## Results and discussion

### Screening

Fifty-three filamentous fungal isolates out of 55 isolates (96.36%) showed a positive reaction for the production of L-asparaginase enzyme as seen due to the color change in a zone surrounding the colonies. Only 2 isolates (2/55 3.63%) (*Penicillium. islandicum* P7 and *Phoma* sp.) didn’t show any change of color of any used indicator. Two isolates of *Eurotium amstelodami* didn’t show any color change in phenol red and bromothymo_l_ containing media but showed a slight color change of zone in the cresol red medium.

### Semi-quantitative screening

In semi-quantitative screening, we found that *P. camembertii* (3.18 EAR) (Figure 1-a),, *C. tenuissimum* (3.66 EAR) (Figure 1-b) and *A. carneus* (3.18 EAR) (Figure 1-c) were the best L-asparaginase strain producer among the tested isolates. Also our results showed that bromothymol blue showed a stronger colour contrast and larger EAR compared with phenol red and cresol red. This result ties well with previous study carried by Mahajan et al. [11] Wherein bromothymol blue (BTB) used instead of phenol red as pH indicator in L-asparagine-containing medium to distinguish between L-asparagine producers and non-producers fungal isolates, _L_-asparagine producers formed dark blue zone around Fungal colonies (medium supplemented with BTB at acidic pH are yellow and it changes to blue at alkaline pH). These authors concluded that BTB is more sensitive and accurate. However, this result contrary to the findings of Mihooliya *et al*. [6] who showed that zone of hydrolysis produced by L-asparagine producing bacteria in media supplemented with cresol red is efficient, better differentiable and consistent contrary to phenol red and BTB, we speculate that this might be due to the use of _L_-asparagine producers bacterial isolates by Mihooliya et al. [11] instead of fungal isolates.

**Figure 1a.**
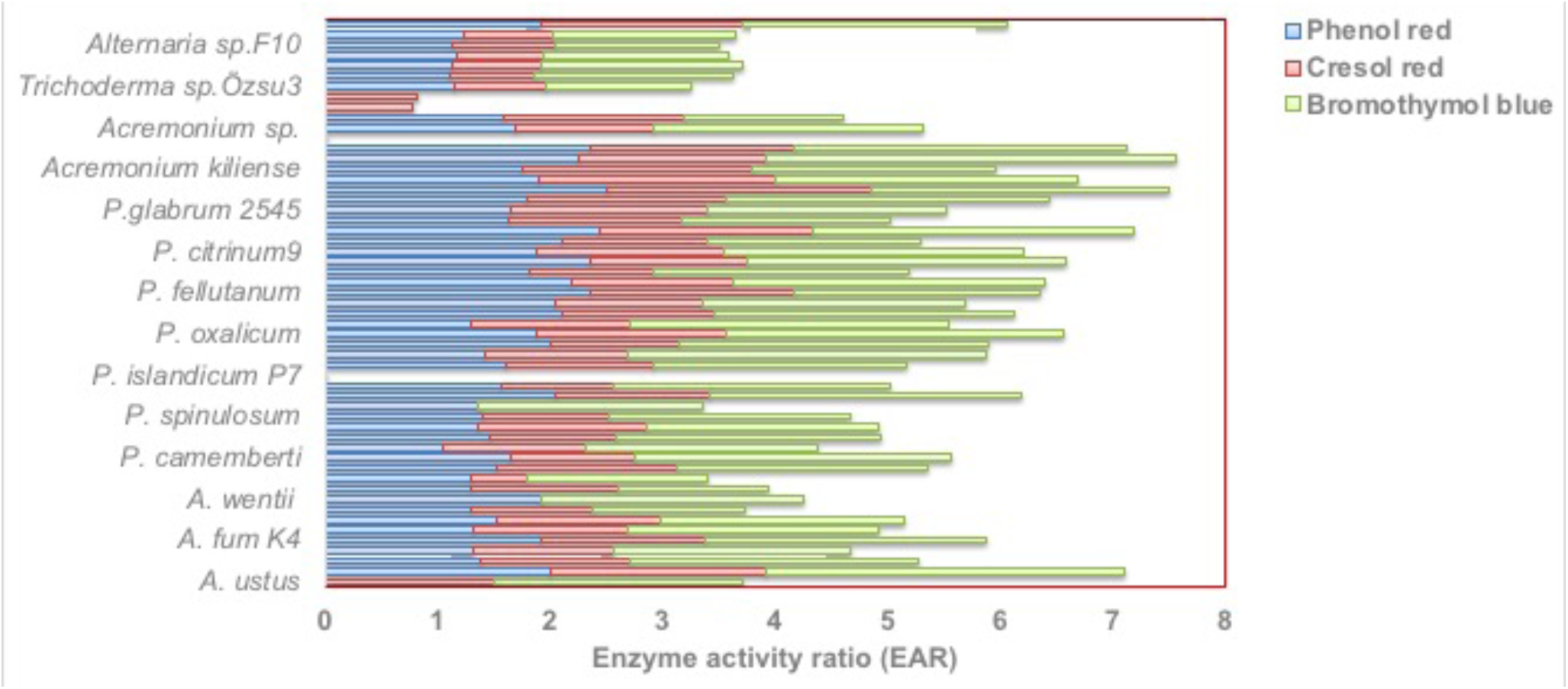
Semi-quantitative screening of different filamentous fungi isolated from different habitats for the production of L-asparaginase with three indicator dyes phenol red, cresol red and bromothymol

**Figure 1-b.**
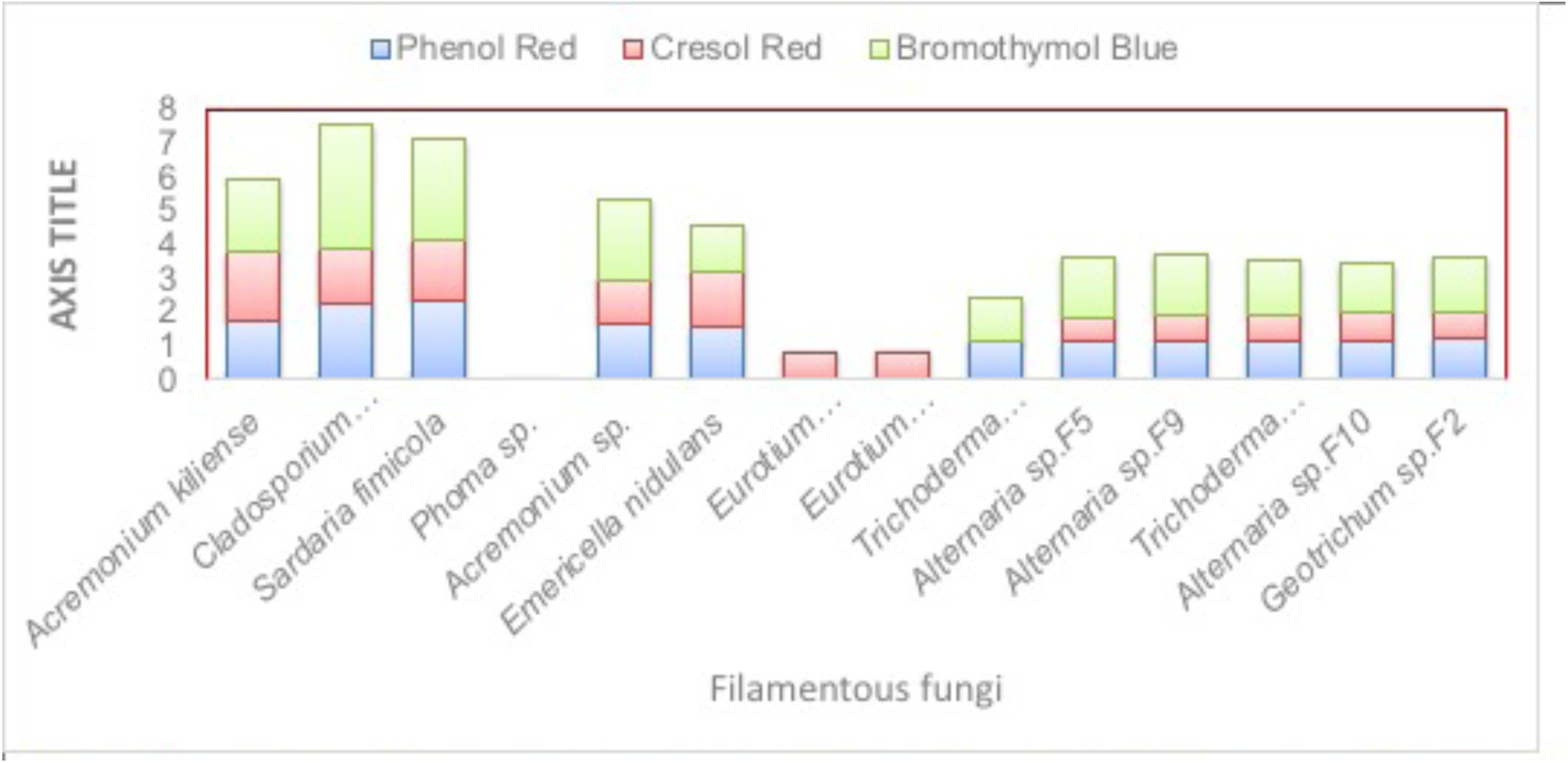
screening of filamentous fungi isolated from different habitats for the production of Lasparaginase with three indicator dyes phenol red. cresol red and bromothymol

**Figure 1C.**
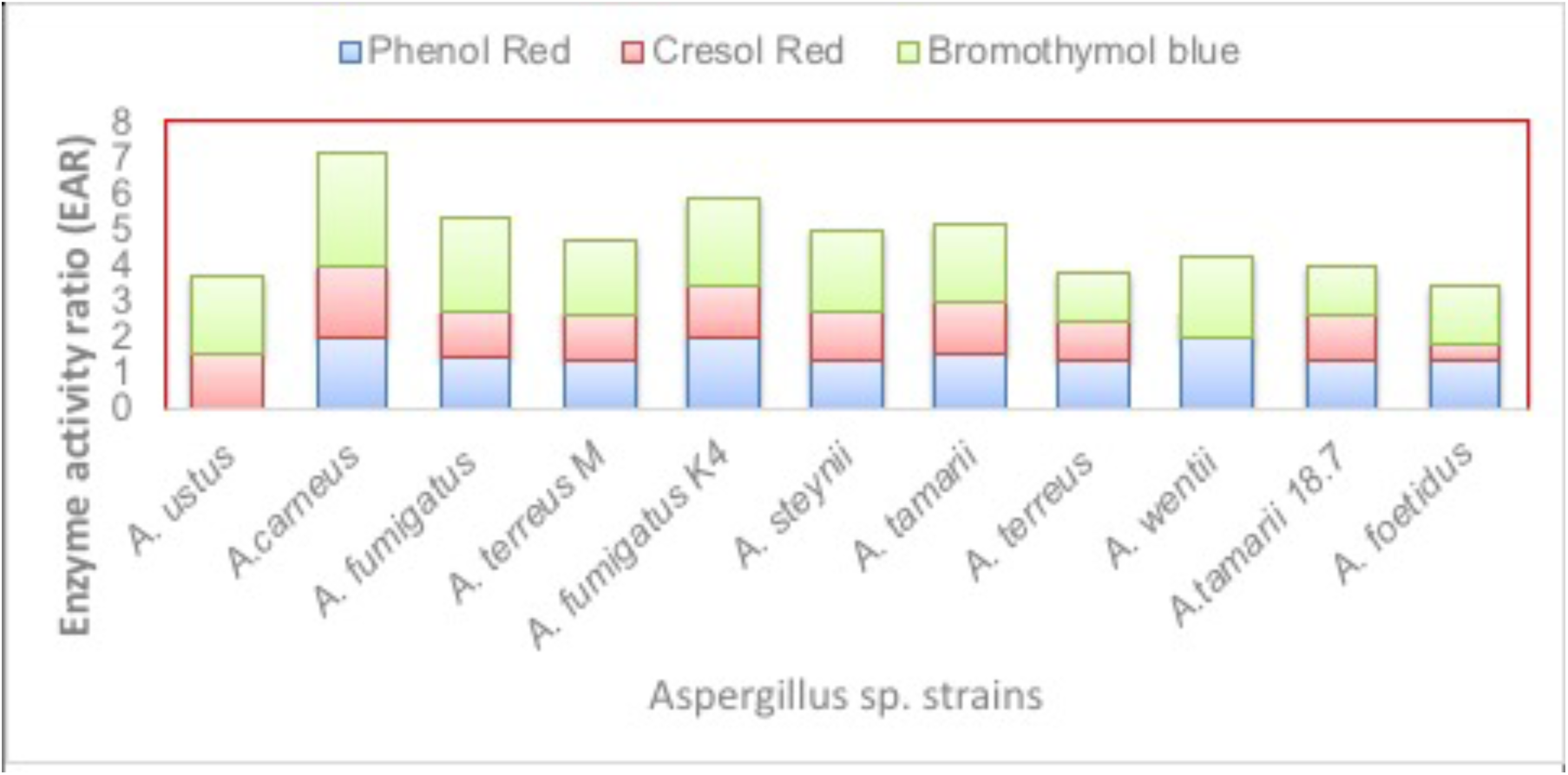
Screening of Aspergillus sp. strains isolated from different habitats for the production of L-asparaginase with three indicator dyes phenol red, cresol red and bromothymol

### Effect of carbon source

Our results showed that all the isolates produced different _L_-asparaginase levels in MCD medium containing 0.2% carbon sources. The highest _L-_asparaginase levels (5.558 U/ml), (4.95 U/ml) and (4.923 U/ml) were produced by *C. tenuissimum, A. carneus* and *P. camemberti*, respectively in MCD medium containing 0.2% glucose (Figure 2). These findings are consistent with research done by Gurunathan and Sahadevan [12], Raja *et al*.[13] and Zia et al. [14].

**Figure 2.**
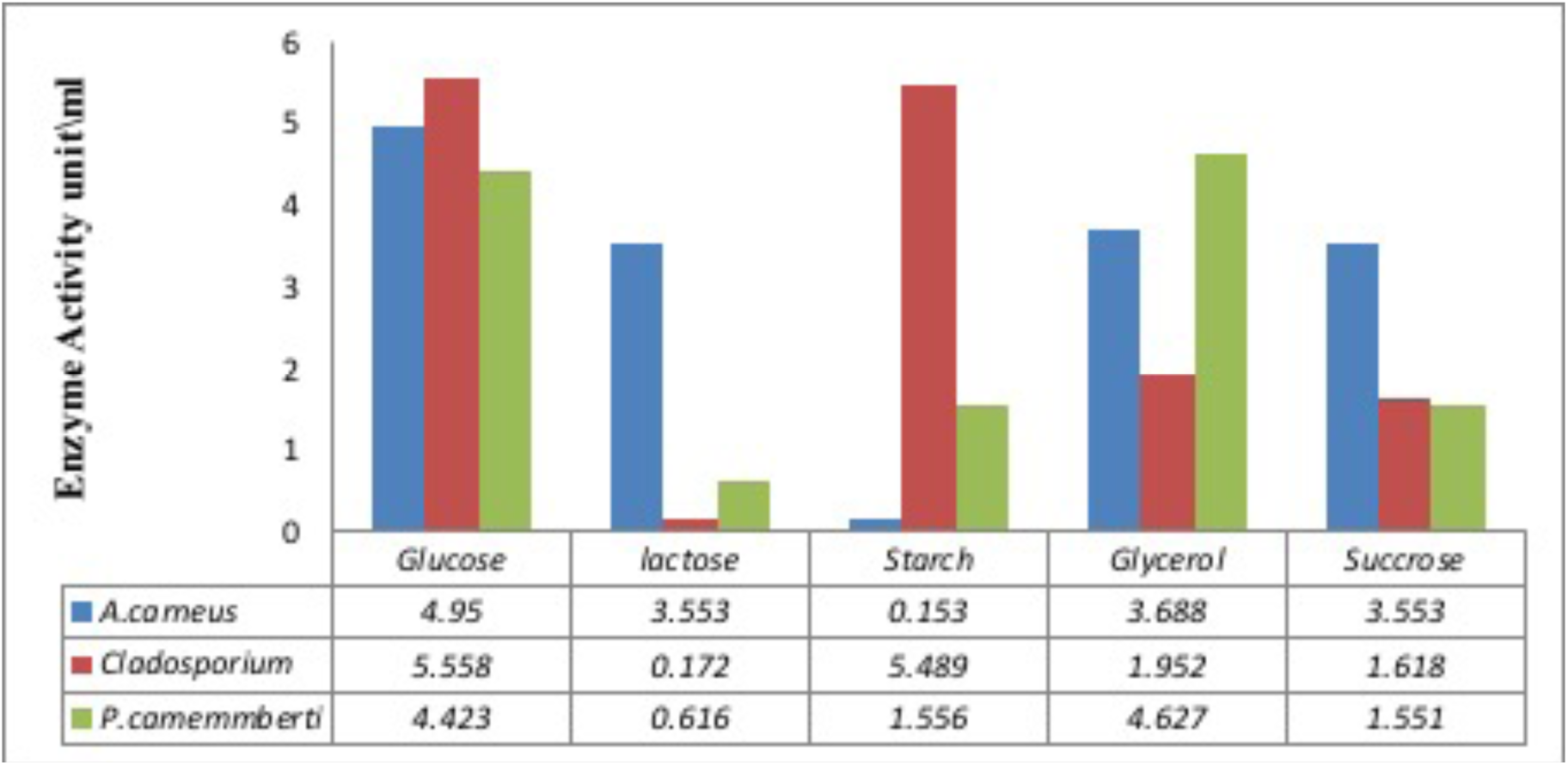
Impact of carbon sources on L-asparaginase production using three different fungal species.

### Effect of nitrogen sources

Also our results demonstrated that there was a high degree of variation regarding the level of produced _L-_asparaginase when changing the type of nitrogen source in MCD medium. *A. carneus* and *P. camemmberti* showed their maximum level of _L-_asparaginase (4.884 U/ml, 3.483 U/ml), respectively when cultivated in MCD media supplemented with ammonium chloride. Moreover, in this study we found that using urea as nitrogen source produced the highest level of _L-_asparaginase (5.015 U/ml) by *C. temimmisium* (Figure 3). The effect of supplementation of ammonium chloride, urea and sodium nitrate as nitrogen sources was studied on growth and production of _L-_asparaginase by *A. terreus*. Urea was identified as the best supplementary nitrogen source [10]. Abbas Ahmed et al. [16] and Sarquis et al. [17] have shown that urea had lowest production of _L-_asparaginase.

**Figure 3.**
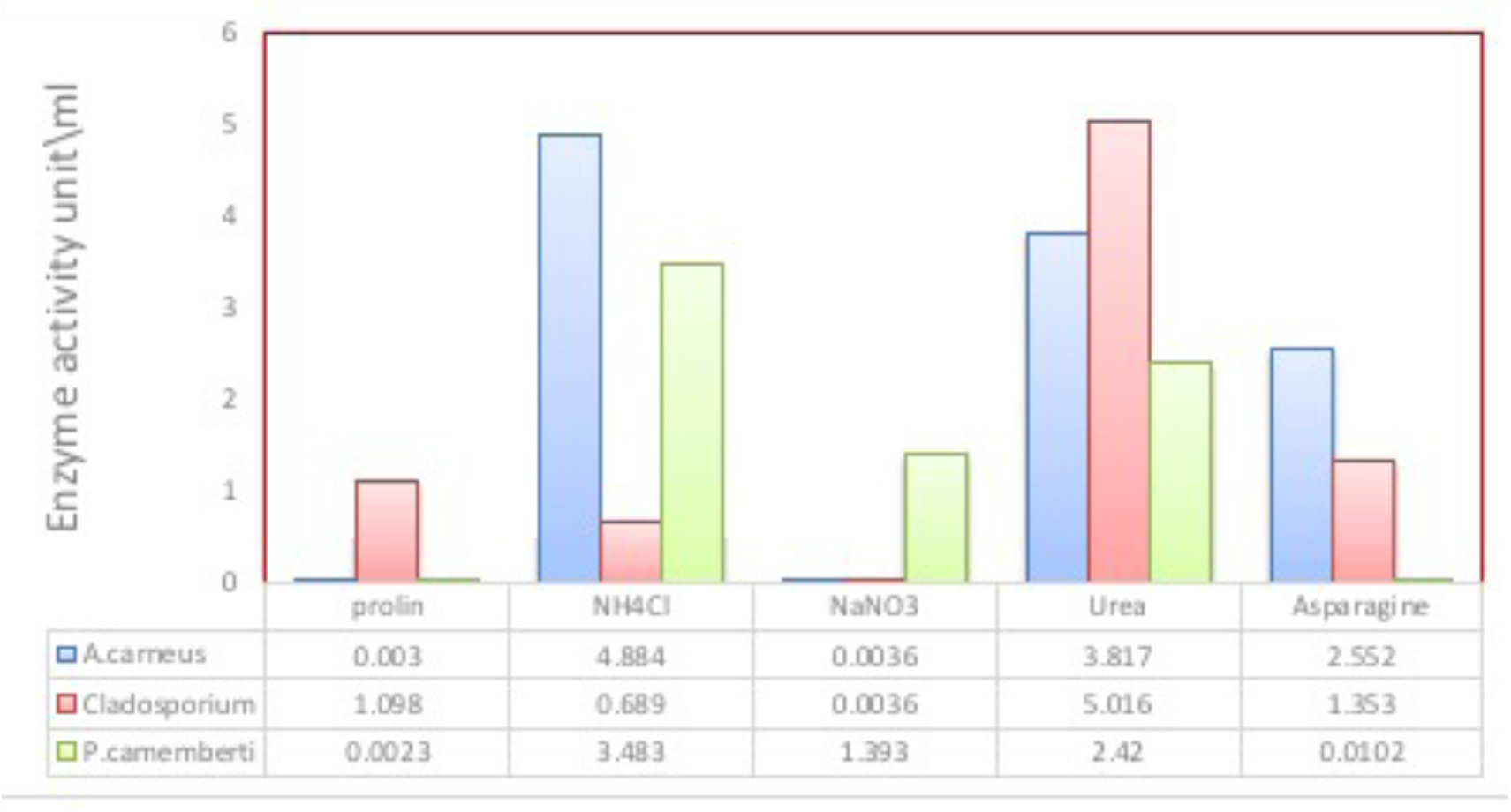
Impact of nitrogen sources on L-asparaginase production using three different fungal isolates.

Generally, proline was a poor substrate, isolates reached their lowest LA levels in a proline-containing medium (Figure 5) which is contradicting the result of Sarquis et al. [17]. The contradiction probably might be due to the difference in the used organism for production.

## Conclusion

This study showed that Penicillium and Aspergillus species can produce L-asparaginase at different levels. Bromothymol blue is the best indicator for screening fungal LA producer. *P. camemberti, A. carneus*, and *C. tenuissimum* are good LA producing isolates. The carbon/nitrogen sources present in the production environment can affect the level of LA produced by the fungi. Glucose is the best carbon source whereas urea was the best nitrogen source for LA production.

